# Arabidopsis species deploy distinct strategies to cope with drought stress

**DOI:** 10.1101/341859

**Authors:** M. Bouzid, F. He, G. Schmitz, R.E. Häusler, A.P.M. Weber, T. Mettler-Altmann, J. de Meaux

**Author notes:** Author for correspondence: Juliette de Meaux.

## Abstract

- **Background and Aims** Water limitation is an important determinant of the distribution, abundance and diversity of plant species. Yet, little is known about how the response to limiting water supply changes among closely related plant species with distinct ecological preferences. Comparison of the model annual species *A. thaliana* to its close perennial relatives *A. lyrata* and *A. halleri*, can help disentangle the molecular and physiological changes contributing to tolerance and avoidance mechanisms, because these species must maintain tolerance and avoidance mechanisms to increase long-term survival, but they are exposed to different levels of water stress and competition in their natural habitat.
- **Methods** We conducted a dry-down experiment that mimics a period of missing precipitation. We quantified the covariation of progressive decrease in soil water content (SWC) with various physiological and morphological plant traits across a set of representative genotypes in *Arabidopsis thaliana, A. lyrata* and *A. halleri.* To quantify the degree of plant stress, transcriptome changes were also monitored.
- **Key Results** The analysis of trait co-variation demonstrates that the three species differ in the strategies they deploy to respond to drought stress. *A. thaliana* showed drought avoidance reaction but failed to survive wilting. *A. lyrata* efficiently combined avoidance and tolerance mechanisms. By contrast, *A. halleri* showed some degree of tolerance to wilting but it did not seem to protect itself from the stress imposed by drought. Transcriptome data collected just before plant wilting and after recovery corroborated the phenotypic analysis, with *A. lyrata* and *A. halleri* showing a stronger activation of recovery- and stress-related genes, respectively.
- **Conclusions** We conclude that these three *Arabidopsis* species have evolved distinct strategies to face drought stress, and discuss the extent to which these strategic differences reflect their respective ecological priorities.

## INTRODUCTION

All physiological and cellular plant aspects depend on water, so limitation in its supply is a major abiotic stress restricting plant growth and crop yield (Stebbins, 1952; Boyer, 1982; Bohnert *et al.*, 1995; Bray, 1997, Lambers *et al.*, 1998; Bray *et al.*, 2000). Water limitation is also a crucial determinant of the distribution, abundance and diversity of plant species (Hoffmann & Sgró, 2011).

All spermatophytes possess the molecular toolkit to tolerate intense cellular dehydration in seeds (Golovina *et al.*, 1997; Kermode, 1997; Wehmeyer & Vierling, 2000). Adult plants can draw from this toolbox to tolerate a certain degree of dehydration in vegetative organs (Ludlow, 1989; Shinozaki & Yamaguchi-Shinozaki, 2007). This tolerance strategy relies on osmotic adjustment via the accumulation of an array of solutes, such as amino-acids, sugars, or dehydrins (Close, 1996). The expression of heat shock proteins, chaperones, or late embryogenesis abundant (LEA) proteins can further help to protect the cell against damages imposed by low internal water potential (Ingram & Bartels, 1996; Reddy *et al.*, 2004,Yue *et al.*, 2006; Szabados, 2010).

However, plants have evolved additional strategies to handle drought stress: escape and avoidance (Ludlow, 1989; Fukai & Cooper, 1995; Verslues & Juenger, 2011; Fang & Xiong, 2015). The escape strategy is based on the adjustment of developmental transitions to elude direct exposure to drought. With an increase in the duration of seed dormancy or a shortening of the life cycle, the plant is simply not facing dry seasons (Fox, 1990; Bewley, 1997; Tonsor *et al.*, 2005; Franks *et al.*, 2007; Kronholm *et al.*, 2012; Lovell *et al.*, 2013). The avoidance strategy, instead, seeks to maintain water levels within tissues through a reduction of water loss and the enhancement of water uptake, so that the plant bypasses the damaging effects of drought (Levitt, 1980; Ludlow, 1989; Price *et al.*, 2002; Farooq *et al.*, 2009; Munemasa *et al.*, 2015).

The relative importance of strategies to cope with drought stress is expected to be intimately linked to the life history and ecology of species. Indeed, tolerance, avoidance, and escape strategies are not independent in evolution (Grime, 1977). Trade-offs between growth and tolerance can constrain their optimization (McKay *et al.*, 2003,Steven, 2011). Annual species prioritize the escape strategy, which in turn can release the need for tolerance mechanisms (Kooyers, 2015). Perennial species, by contrast, must maintain tolerance mechanisms to increase long-term survival.

Dehydration triggers dramatic responses in plant cells, as indicated by the fast and extensive changes in gene transcript levels (Shinozaki & Yamaguchi Shinozaki, 2000; Iuchi *et al.*, 2001; Seki *et al.*, 2001; Shinozaki & Yamaguchi, 2007; Matsui *et al.*, 2008; Harb *et al.*, 2010). Part of this response is regulated by the key drought-stress hormone abscisic acid (ABA), but ABA-independent transcriptional regulation also plays an important role (Iuchi *et al.*, 2001; Seki *et al.*, 2001; Sakuma *et al.*, 2006; Yoshida et *al.*, 2014; Urano *et al.*, 2017). The complex architecture of gene regulatory responses to stress is believed to contribute to restricting the reactions at cell and whole-plant levels when the internal water potential drops (Bray, 1997; Szabados, 2010; Osakabe *et al.*, 2014). By articulating growth and stress responses, transcriptomic changes take part in both the deployment of avoidance strategies and the promotion of recovery from stress, yet they also reveal the degree of stress sensed by the organisms. Distantly related annual species, such as rice and *Arabidopsis*, show common patterns of stress responses (Nakashima *et al.*, 2009). Much less is known about how responses to stress are reshaped in closely related species with strongly divergent ecologies and life-histories.

Comparison of *A. thaliana* to its close relatives can help disentangle the molecular changes contributing to tolerance and avoidance mechanisms, because different species in the genus have evolved distinct ecologies with contrasting demands on tolerance and avoidance (Clauss & Koch, 2006). The model species *A. thaliana* shows a broad distribution range from north of Scandinavia to Africa (Hoffmann, 2005,Durvasula *et al.*, 2017). Its response to severe or mild drought stress has been described in detail (Seki *et al.*, 2002; Bray, 2004; Verslues & Juenger, 2011; Des Marais *et al.*, 2012; Juenger, 2013; Bechtold *et al.*, 2015; Lovell *et al.*, 2015). Several studies point to the adaptive relevance of its variation (Kesari *et al.*, 2012; Exposito-Alonso *et al.*, 2017). This annual species can also rely on modifications of its life cycle to adjust the timing of escape and/or avoidance strategies to drought threats (McKay *et al.*, 2003; Kronholm *et al.*, 2012; Wolfe & Tonsor, 2014). The two sister species *Arabidopsis lyrata* and *A. halleri*, by contrast, are less likely to rely on escape strategies because year-to-year survival is of major importance for these perennials. *A. lyrata* is probably the most exposed of the two to natural selection by drought due to its preference for low competitive communities in soils that do not retain water (Clauss & Koch, 2006; Ellenberg & Leuschner, 2010; Sletvold & Agren, 2012). *A. halleri*, instead, must grow to out-compete other species in crowded habitats (Clauss & Koch, 2006; Ellenberg & Leuschner, 2010; Stein *et al.*, 2017). Its specific ability to accumulate heavy metals enhances its defenses against herbivores but sets strong constitutive demands on detoxifying systems which are important for reestablishing homeostasis after stress (Mittler, 2002; Becher *et al.*, 2004; Krämer & Clemens, 2006; Stolpe *et al.*, 2016). The contrasted ecologies of these three species thus predict major consequences on their strategies to face up with the challenges imposed by water limitations.

To test this prediction, we set up an experiment to infer the response strategy to drought of sets of accessions representative of the three species *A. thaliana*, *A. halleri* and *A. lyrata*. For this, we measured plant drought reaction at both phenotypic and transcriptomic levels in a dry-down experiment that mimics the progression of water depletion in natural conditions. Our data showed that species deploy different avoidance and tolerance strategies in response to decreasing levels of <u>s</u>oil <u>w</u>ater <u>c</u>ontent (SWC).

## MATERIALS AND METHODS

### Plant material and growth conditions

Altogether, 16 to 22 and 12 to 17 central European *A. lyrata* and *A. halleri* accessions, respectively, were included in the dry down experiments. The accessions were taken from populations representative of the diversity described in these species (Supplementary Table S1, Pauwels *et al.*, 2005; Ross-Ibarra *et al.*, 2008; Novikova *et al.*, 2016; Stein *et al.*, 2017). They were compared to 16 *A. thaliana* accessions from Spain with European genomic background (The 1001 Genomes Consortium 2016). This sample was chosen because i) the populations are among the most drought resistant in *A. thaliana* (Exposito-Alonso *et al.*, 2017) and ii) are late flowering (Arapheno database, FT16, DOI: <u>0.21958/phenotype:262</u>) so that the stress exposure cannot be circumvented by life cycle termination. For each accession, five replicates (vegetatively propagated clones for the self-incompatible species, single-descent seeds for *A. thaliana*) were distributed in 5 randomized complete blocks.

Plants were grown in 7×7×8 cm pots filled with 150 g of a well-homogenized mixture of VM soil (60 to 70% of peat and 30 to 40% of clay), perlite and seramis (clay granules) in a CLF controlled growth chamber (Perkin Elmer, USA). Growth conditions were 10 h (20°C): 14 h (16°C), light: dark, at a photon flux density (PFD) of 100 μmol m^-2^ s^-1^ supplemented with 10 min of dark-red light at the end of the day. Relative humidity was set to 60%.

### Dry-down experimental design

Plants were grown for five weeks in the greenhouse, re-potted in weighed pots filled with the initial soil mixture, and transferred to the growth chamber. Soil moisture was quantified every day (*X_t_*) by monitoring pot mass with a precision balance with an accuracy of 0.01 g. To calculate the soil moisture, several pots were fully dried down in an oven to estimate the weight of dry soil (*X_0_*) in the initial soil mixture and subsequently saturated with water to determine the weight of 100% wet soil (*X_f_*). The percentage of soil moisture was calculated as [*(X_t_ - X_0_)* / (*X_f_* - *X_0_*)] × 100. For acclimation, plants were grown for two weeks in pots with 60% soil moisture. After acclimation, plants were not watered until showing first symptoms of wilting. Plants were re-watered two days after wilting. One to two weeks later survival and symptoms of damage were scored.

Three independent biological experiments were performed. We discarded any plant that was not healthy and vigorously growing at the start of the experiment. Focusing on initially healthy plants thus resulted in slight differences in the number of replicates and/or accessions (for details see Supplementary Table S1-S3). The two first experiments were used for phenotypic characterization and the third for sampling of leaf material for RNA extraction. In the experiment, plants were re-watered on the day of wilting to allow collecting leaf material after recovery.

### Phenotypic trait measurements

#### Phenotypic differences between species in well-watered conditions

Three phenotypes were measured in *A. halleri* and *A. lyrata* in glasshouse-grown plants under well-watered conditions: stomatal density, stomata length, and carbon isotope discrimination (δ^13^C). Stomatal density and length were quantified in fully-developed leaves of five replicates of nine accessions per species following protocol described by Paccard *et al.*, (2014). δ^13^C in one fully developed leaf was quantified for 4 replicates of the same nine accessions of each species according to the method used by Gowik *et al.*, (2011).

#### Phenotypic variation in response to soil dry-down

Eight phenotypes were measured during the dry-down experiment. Rosette leaf area was quantified on day zero of the dry-down experiment, using ImageJ to separate green pixels from the background images and RosetteTracker (Vylder *et al.*, 2012) to convert total green pixel into mm^2^. The first day we observed that leaves had lost their turgidity was scored as wilting day. Soil moisture was measured every day until the day of wilting. The rate of soil water loss was calculated for each pot over the first seven days after water withdrawal. Leaf lamina thickness was measured on one ink-marked medium-size leaf every second day using a digital ruler (HOLEX, Hoffmann Group, Knoxville, USA) with an accuracy of 0.03 mm. Efficiency of the photosynthetic light reaction was measured by Pulse-Amplitude-Modulation (PAM) fluorometry (Schreiber *et al*., 1986) using the IMAGING-PAM-Series (M-Series-Maxi version, Heinz Walz GmbH, Effeltrich,Germany). In order to gain information on the intactness of photosystem II (PSII) and hence its potential photosynthetic capacity, the maximum quantum efficiency of open PSII reaction centers (F_v_: F_m_, i.e. the ratio of variable to maximum Chl*a* fluorescence) was determined (Genty *et al.*, 1989; Maxwell & Johnson, 2000). Before the application of a saturating light flash (duration 0.8 s), plants were dark-adapted for 30 min. Intact and non-stressed plants usually show an F_v_: F_m_ ratio of around 0.8. Plants that developed new leaves within two weeks after re-watering were scored as having survived and the damage caused by wilting was quantified visually on a damage severity scale from one to five, reflecting the percentage of damaged leaf area, leaf color and leaf strength. The number of days of tolerated wilting was scored on plants that survived the first dry-down experiment. For this, plants were dried down a second time until wilting and re-watered after three, four, five, or six days of wilting. Despite previous exposure to drought stress, plants wilted at the same limiting SWC (e.g. approximately 20%), suggesting that if plant show differences in stress memory, this effect is not detectable after 3 weeks. Photosynthetic activity and duration of tolerated wilting were measured in the first experiment, whereas rosette area and leaf thickness were measured only in the second experiment (Supplementary Table S2).

### Statistical analysis of phenotypic variation

All plots were created using the *CRAN-package ggplot2* (Wickham, 2009). We used generalized linear models (R function glm) and multiple comparison tests using the *Simultaneous Inference in General Parametric Models* package named *multcomp* and Tukey’s Honest Significant Difference test (Tukey HSD). For each phenotype, we ran several models. As we did not detect any block effect for the different measured traits, we removed it from our models. Following are the different tested models, and later in the results part, we will mention which was the best model:

(M1) tests the accessions nested within species effect
Y_ijk_=μ + α_i_ species + β_ij_ (species _i_ accession _j_) + ϵ_ijk_
(M2) tests only the species effect when the accession effect is not significant
Y_ij_=μ + α_i_ species _i_ +ϵ_ij_
(M3) tests the interaction between species and time effect
Y_ijk_= μ+ α_i_ species _i_ + β_j_ time _j_ + γ _ij_ (species _i_ time _j_) +ϵ_ijk_
(M4) tests the effect of interaction between species and the cofactor of interest
Y_ijk_=μ + α_i_ species _i_ + β_j_ cofactor _j_ + γ_ij_ (species _j_ cofactor _j_) +ϵ_ijk_

Where:

Y: quantitative dependent variable e.g. measured phenotypic trait; μ: is the overall mean; α, β and γ regression coefficients; species; accession; time; cofactor (e.g. initial rosette size, desiccation rate, initial leaf thickness, damage scores, days after wilting etc.): independent variables with the different levels i, j, and k; ε prediction error.

Different error distributions were specified for each phenotypic trait, depending on whether or not overdispersion was detected (i.e. whether the residual deviance was of the order of magnitude of the degrees of freedom). A negative binomial fitted best the number of days until wilting, soil moisture, initial rosette area, initial leaf thickness, damage scores, relative leaf water loss, stomatal density and stomata length. A Gaussian distribution fitted better measures of desiccation rate and δ^13^C, a quasi-Poisson distribution was used for the photosynthesis activity and quasi binomial distribution for survival rate. We performed an ANOVA using Fisher’s test (or Chi test for the binomial distribution of error) to identify the best model (P-value ≤ 0.05).

### Analysis of transcriptome variation during dry-down

In the third dry-down experiment, three to four young leaves of ‘hal2.2’ and ‘Plech61.2a’, typical accessions of *A. halleri* and *A. lyrata*, respectively, were sampled from three replicate individuals at three time points: 1) before water withdrawal (soil moisture around 60%), 2) before wilting symptoms appeared (20% to 25% of soil moisture), and 3) leaves formed during the recovery phase (10-15 days after re-watering). These two accessions are representative of the phenotypic diversity observed in the dry-down experiment. RNA extraction was performed using the *PureLink™ RNA Ambion Mini Kit* (Thermofisher, Darmstadt, Germany). RNA quality and quantity were checked by Agilent 2100 bioanalyzer (Agilent Technologies, Palo Alto, Calif.) using RNA nano chips. RNA of 18 leaf samples was sequenced on Illumina *HiSeq4000* by the Cologne Center for Genomics. Raw sequence data are available in the SRA database under the accession number: SRP150056.

We used the *fastx-tool-kits* from the *FastQC* package (V0.11.4) for raw sequence quality trimming and filtering following He et al. (2016). Low quality nucleotides were removed from the 3′-ends of the sequences using 20 as a phred score threshold (t) and 50 as minimum length (l). Sequences were reverse complemented using fastx_reverse_complement to cut the other end as we did for the 3′-end. Reads with less than 90% bases above the quality threshold and paired-end reads with a single valid end were discarded. We used the software package *STAR* with standard parameters (Dobin & Gingeras, 2015) to map trimmed and filtered reads to the *A. lyrata* reference genome V1 (Hu *et al.*, 2011). Alternative transcripts were not considered because the current annotation of the *A. lyrata* genome does not describe alternative transcripts. Transcriptome sequencing yielded a total of 15 million read pairs per sample, with a read length of 75 bp. We used ‘samtools view –q 10’ to select the uniquely and high quality mapping reads with a probability of correct mapping of 90%.

On average, more than 80% of all reads were uniquely mapped and around 20% of unmapped and multiple mapped reads (Supplementary Fig. S1). R scripts were used to verify that reads covered the whole length of genes (and confirm that we had no sign of RNA degradation) and for counting the number of reads mapped to each. The *DESeq2* Bioconductor package from *R (Bioconductor version: Release 3.5)* was used to find genes that were differentially expressed (DE) between the different conditions (Love *et al.*, 2014). We used the Wald test to compute P values and the following design: ~ species/sample point, with two levels for the factor species (*A. halleri* and *A. lyrata*), and three levels for the factor sample point (leaves sampled at 60% of soil moisture, at 20-25% of soil moisture, and after recovery). Genes with a P value < 0.1 after Benjamini-Hochberg correction for false discovery rate (FDR) and log_2_-fold change ≤ −0.5 or ≥0.5 were considered as DE.

### Gene ontology analysis

Functional enrichments among DE genes were performed using *org.At.tair.db* data package of *Bioconductor* and the rank test of the *TopGO* package (Alexa & Rahnenfuhrer, 2010) was used to identify enriched gene ontology terms. The *elim* algorithm followed by a *Fisher* test were used with a cut-off of 0.01. As background all expressed genes were used (around 12220 genes). Enrichments were analyzed separately for: 1) all responsive genes, 2) down-regulated genes, and 3) up-regulated genes. The hyper-geometric test was used to test for the significance of gene overlap with a set of stress responsive genes (Matsui *et al.*, 2008).

## RESULTS

### Interspecific differences in stomatal density and stomata length but not in water-use efficiency

We evaluated whether, under well-watered conditions, constitutive physiological differences between *A. lyrata* and *A. halleri* can influence their potential to face limiting SWC. Variation in stomatal density on the leaf surface was explained by both within and between species variance (M1: F_18, 469_ = 36.15, P-value < 2e^-16^ within species; F_1, 487_=256.59, P-value < 2.2e^-16^, between species, Fig. 1A).

In *A. lyrata* stomatal density on the abaxial leaf surface was lower than in *A. halleri* (on average 80 in *A. lyrata* and 150 stomata mm^-2^ in *A. halleri*). By comparison, a recent and exhaustive analysis of stomatal density in *A. thaliana*, reported that stomatal density varies from 87 to 204 stomata mm^-2^ and it is negatively correlated with stomata length (Dittberner *et al.*, 2018). Stomata were larger in *A. lyrata* compared to *A. halleri* (M1: P-value< 2e^-16^) and the genetic variation in stomata length was significant both within and between these two species (M1: F_16, 1370_ = 53.68, P-value < 2e^-16^ within species; F_1, 1386_=3801.39, P-value < 2.2e^-16^, between species). These differences however did not coincide with differences in carbon isotope discrimination (δ^13^C),a commonly used proxy for water-use efficiency (WUE, Farquhar & Richards, 1984; Farquhar *et al.*, 1989; Lambers *et al.*, 1998; Dawson *et al.*, 2002). In non-stressed conditions, leaf δ^13^*C* showed significant genetic variation within species, but not between *A. halleri* and *A. lyrata* (−29.38 ‰ in *A. lyrata* and −29.37 ‰ in *A. halleri*, on average, M1: F_16, 54_= 7.440, P-value= 9.76e^-09^ within species, and F_1, 70_ = 0.005, P-value =0.969, between species Fig. 1B).

**Figure 1:**
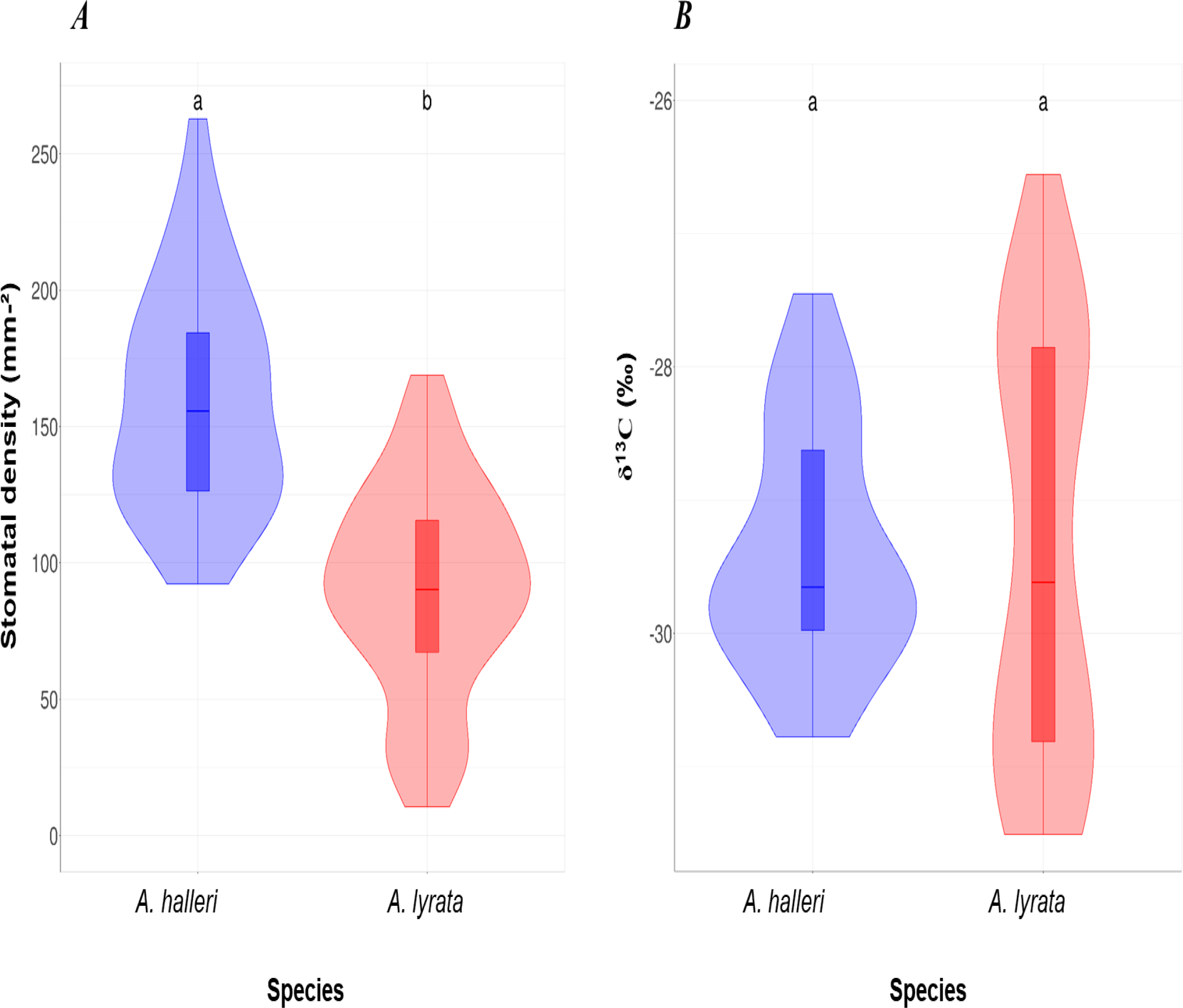
Stomata density and δ^13^C measured in *Arabidopsis halleri* and *A. lyrata* grown under well-watered conditions. (A) Abaxial stomatal density. (B) δ^13^C measured for the same plants. Violin plots with the same letter are not significantly different according to Tukey’s HSD (P value <0.05).

### Wilting-related phenotypes revealed different drought response strategies

The day of first appearance of wilting symptoms differed significantly between species in the first experiment, although accessions within species also differed (M1: F_2, 214_=316.48, P-value < 2.2e^-16^ for species, Fig. 3A, F_48, 166_ =3.51, P-value=1.159e^-09^, for accessions within species). The same result was observed in the second experiment (M1: F_2, 201_= 115.27, P-value < 2.2e^-16^, F_33, 168_= 1.97, P-value= 0.002, Supplementary Fig. S2A).

Wilting manifested differently in the three species. In *A. thaliana*, leaves became pale and curled laterally, in *A. lyrata*, they curled apically, and in *A. halleri* leaves changed to darker green and collapsed (Fig. 2). On average, *A. halleri* accessions wilted around five to seven days after water withdrawal, *A. lyrata* accessions after 12 days and *A. thaliana* accessions after 18 days (Fig. 3A, Supplementary Table S4). Differences in the timing of wilting did not exactly coincide with SWC differences. At wilting, *A. halleri* and *A. lyrata* showed similar soil moisture (18-20%), whereas *A. thaliana* only wilted after soil moisture dropped below 10% (Fig. 3B, Supplementary Table S5). Again, these effects were consistent across experiments (Supplementary Fig. S2B). Significant differences were detected between species for soil moisture at wilting (M1: F_2, 214_ =44.27, P-value=3.982e^-16^, F_2, 201_ =181.60, P-value < 2.2e^-16^ for the first and second experiment respectively), and within species (M1: F_48, 166_ =1.52, P-value=0.020, F_33, 168_ =2.23, P-value=1.07e^-10^ for the first and second experiment respectively).

**Figure 2:**
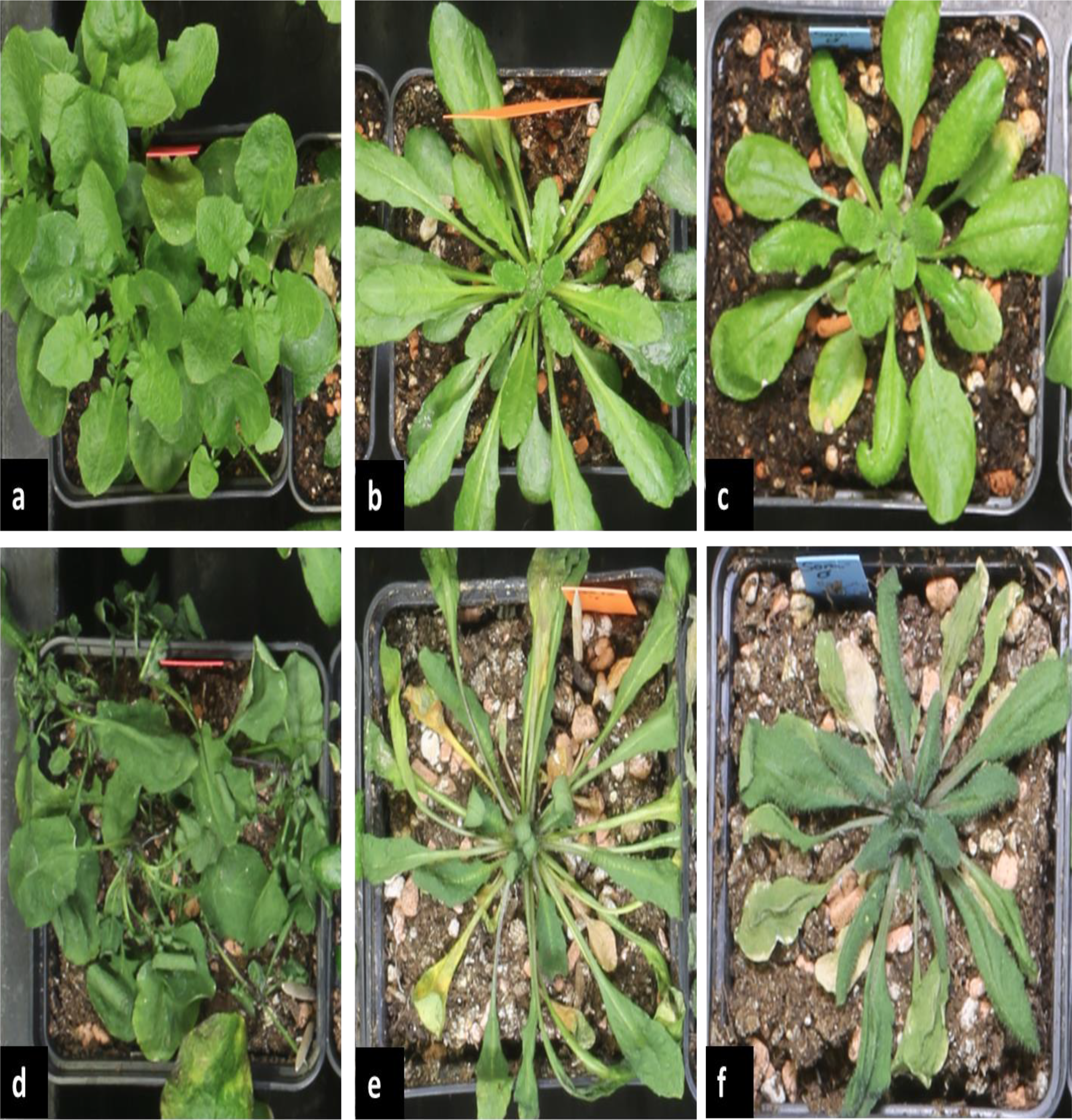
Typical phenotypes of wilting observed in *Arabidopsis halleri*, *A. lyrata*, and *A. thaliana*. Plant morphology before the water withdrawal treatment (top row) and at wilting (bottom row) for A. halleri (a, d), A. lyrata (b, e) and A. thaliana (c, f). All plants were grown in 7cm pots. One single plant was grown in each 7cm pots.

**Figure 3:**
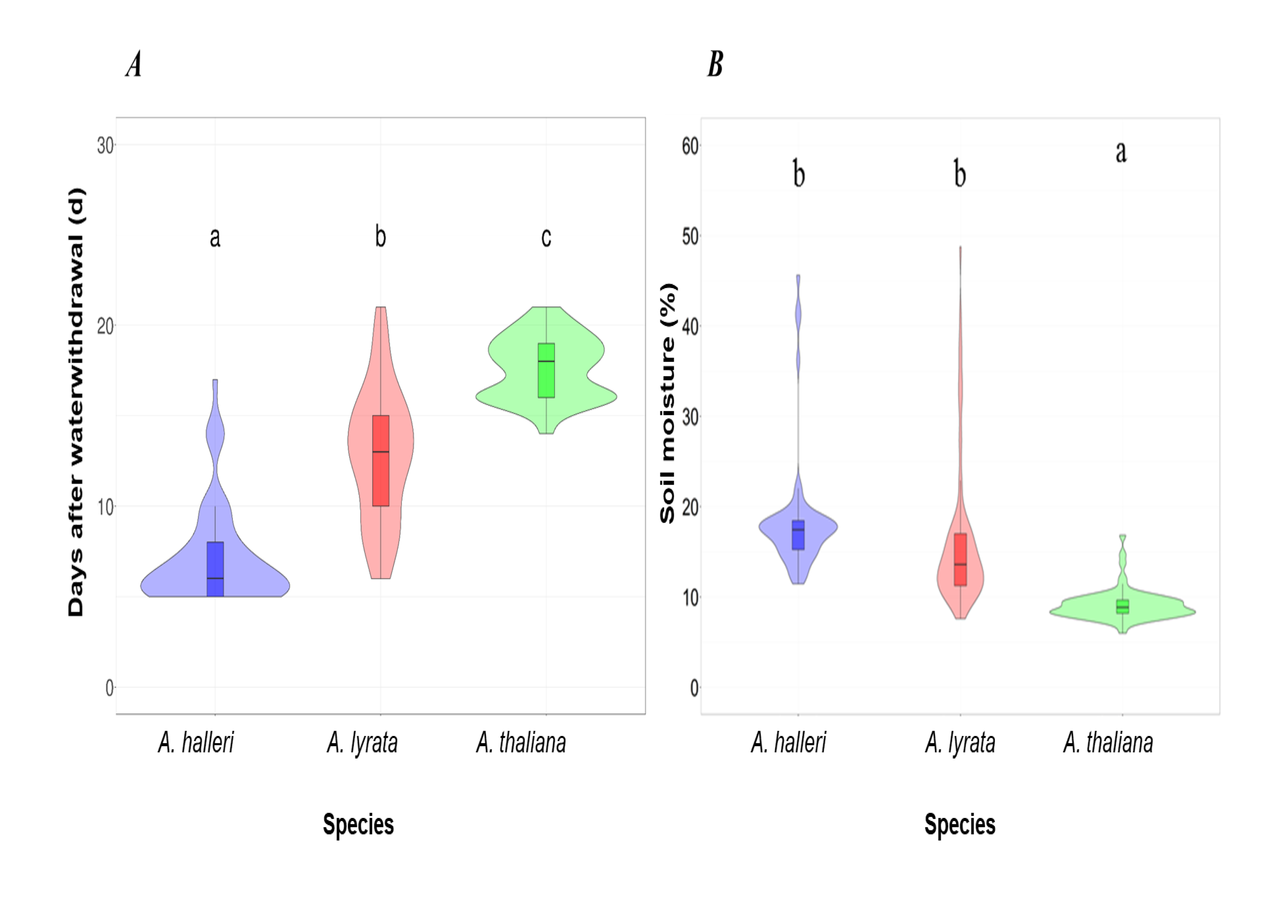
Wilting day and soil moisture at wilting for *Arabidopsis halleri*, *A. lyrata*, and *A. thaliana*. ***(A)*** Number of days between initiation of soil dry-down treatment and wilting. ***(B)*** Soil moisture at wilting. Letters above violin plots indicate significant differences between species (*Tukey’s HSD test, P value <0.05*). Results are shown for the first biological experiment.

### A. halleri plants exhaust SWC faster

To understand why *A. halleri* plants wilted around one week earlier than *A. lyrata* but at a similar soil moisture, we evaluated the rate of soil water loss for each species. We detected a significant interaction between species and time on soil moisture before wilting which showed that soil moisture decreased faster in pots where *A. halleri* accessions grew (Supplementary Fig. S3A, M3: F_12, 1194_ = 97.026, P-value < 2.2e^-16^). *A. halleri* thus consumed water significantly faster than *A. thaliana* and *A. lyrata.* Here again, this observation was replicated in the second biological experiment (M3: F_4, 1224_= 761.07, P-value < 2.2e-16, Supplementary Fig. S3B).

To examine the impact of plant size on the rate of soil water loss, we measured initial plant size and estimated the desiccation rate, defined as the rate of soil water loss per day over the seven days following the water withdrawal in the second experiment of the dry-down experiment. *A. lyrata* and *A. halleri* accessions started with similar rosette size, but *thaliana* rosettes were initially larger (M2: F_2, 173_=10.85, P-value= 3.65e-05, Supplementary Fig. S4A and Table S6). We detected a significant effect of the initial rosette area on the desiccation rate (M4 F_1, 170_=16.10, P-value=8.97e^-05^) but no significant interaction between initial rosette area and species on desiccation rate (M4: F_2, 170_=1.89, P-value=0.15). Therefore, the consumption of soil water does not scale with plant size even though significant correlations between desiccation rate and initial rosette size were detected in *A. halleri*, less in *A. thaliana* but not in *A. lyrata* (Fig. 4A).

**Figure 4:**
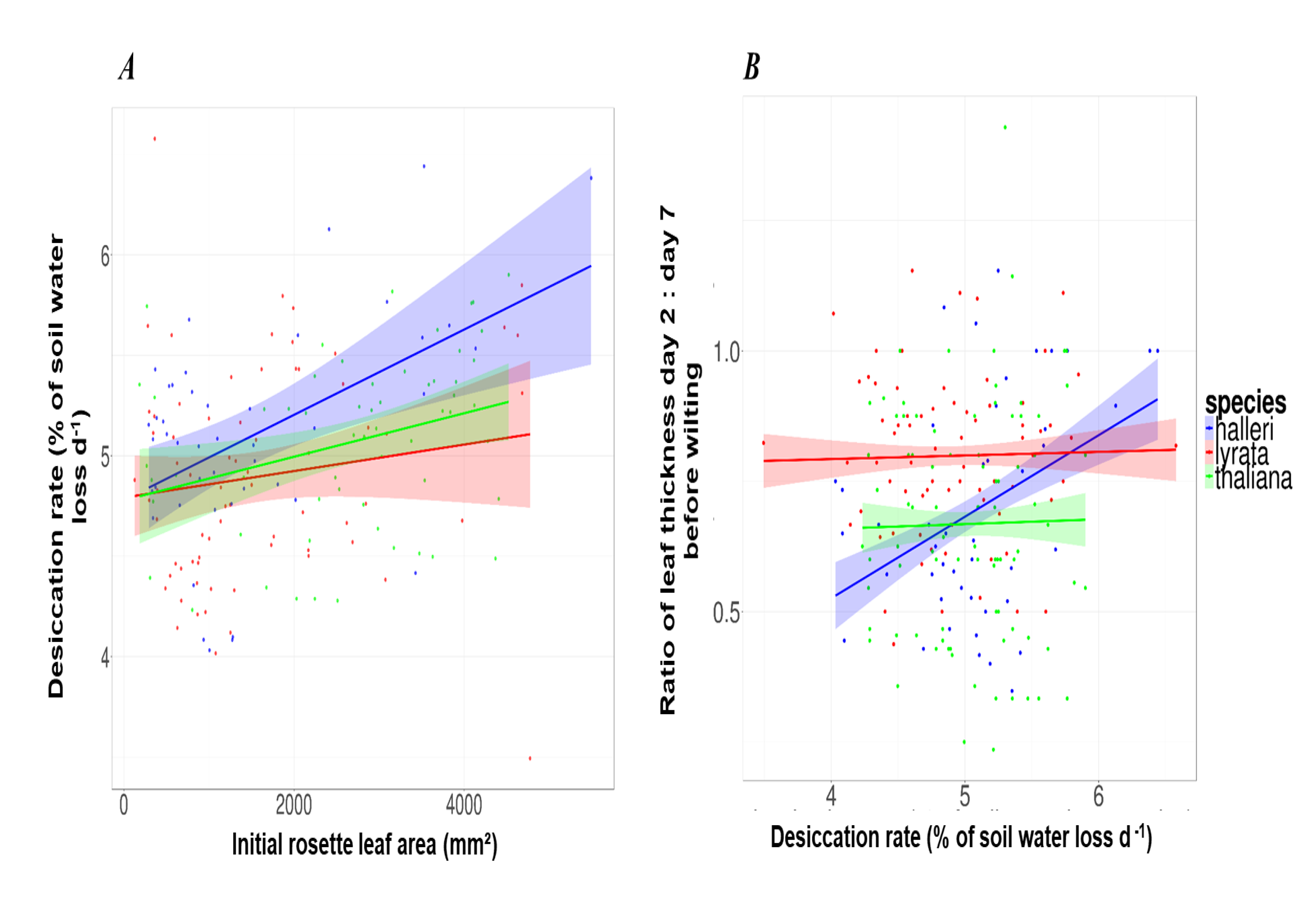
Correlations between desiccation rate and initial leaf size and desiccation rate and the relative leaf water loss. ***(A)*** Correlation between the initial rosette leaf area (at 60% of soil moisture) and the percentage of soil desiccation rate (Pearson correlation coefficients and p values for: *Arabidopsis thaliana* (r = 0.32, P value = 0.013); *A. lyrata* (r = 0.14, P value = 0.22) and *A. halleri* (r = 0.48, P value = 0.00072). ***(B)*** Correlation between the relative water loss in leaves before wilting (equivalent to the ratio of leaf thickness day 2: day 7 before wilting) and the desiccation rate (Pearson correlation coefficients and p values for: *A. thaliana* (r = 0.018, P value = 0.732); *A. lyrata* (r = 0.023, P value = 0.692) and *A. halleri* (r = 0.39, P value = 4.282.10^-08^). Results are shown for the second biological experiment. Lines represent a linear regression smoothing where the shaded ribbons represent the standard error.

### A. lyrata has the lowest relative loss of leaf water content before wilting

To estimate changes in leaf water content during the water-limited phase, we monitored leaf thickness (Lambers *et al.*, 1998) during soil dry-down phase in the second biological experiment. Initial leaf thickness was significantly higher in *A. lyrata* plants compared to *A. thaliana* and *A. halleri* (M1: F_2, 140_=9.38, P-value=3.30e^-10^, Supplementary Fig. S4B and Table S7). We also detected a significant accessions effect within *A. lyrata* on the initial leaf thickness (M1, F_33, 140_= 1.642, P-value=0.02548).

The significant interaction effect of soil desiccation rate and species (M4, F_2, 818_=11.15, P-value=1.66e-05) on leaf thickness change over time revealed that the correlation between leaf thickness and soil desiccation rate was significant only for *A. halleri* (Fig. 4B, Supplementary Table S9). Furthermore, this analysis showed that *A. thaliana* leaves were able to hold higher amounts of water at lower soil moisture, compared to *A. lyrata* and *A. halleri* (Fig. 5), an indication that this species can effectively avoid the effects of drought by maintaining a comparatively higher water content in its leaves.

**Figure 5:**
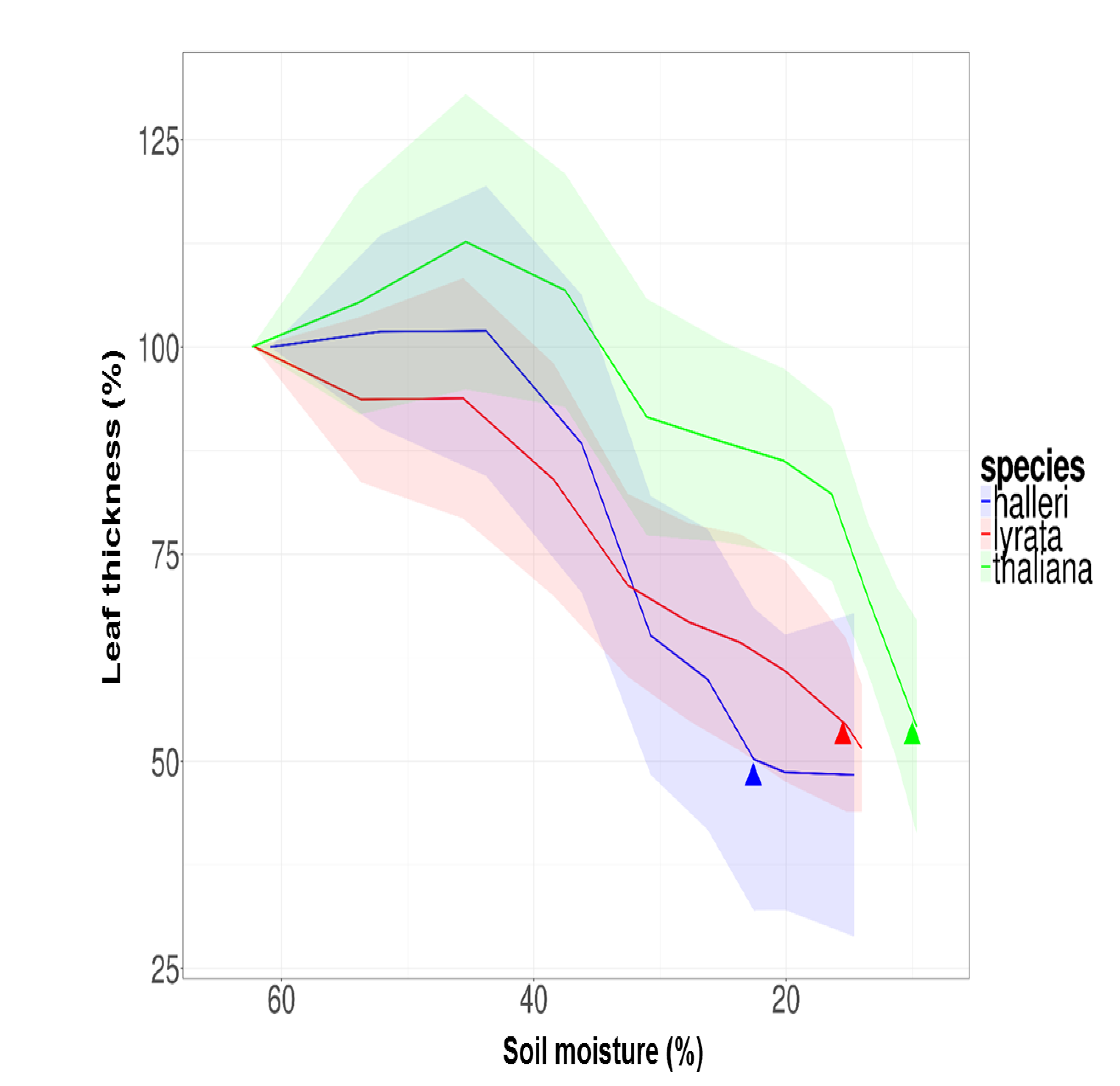
Leaf thickness in response to decrease of soil moisture for *Arabidopsis thaliana*, *A. halleri*, and *A. lyrata*. Results were collected in the second biological experiment. Shaded ribbons represent the standard deviation. Filled triangles correspond to the average wilting soil moisture for the different species.

*A. thaliana* and *A. halleri*, however, lost similar amounts of water in the days preceding wilting. The relative loss of leaf water content before wilting was calculated by the ratio of leaf thickness two days before wilting by leaf thickness seven days before wilting (Fig. 6). There was no significant accessions effect on the decrease of leaf thickness in the seven days before wilting (M1: F_33, 138_= 0.9401, P-value=0.566) but the relative decrease before wilting was significantly higher in *A. thaliana* and *A. halleri*, compared to *A. lyrata* (M1: F_2,171_=6.628, P-value= 5.00e^-8^, Fig. 6, Supplementary Table S8). This pattern indicates that leaf water content in the days preceding the onset of wilting decreased more slowly in *A. lyrata* plants compared to *A. halleri* and *A. thaliana*. This suggests that wilting *A. lyrata* leaves experience lower loss of turgor.

**Figure 6:**
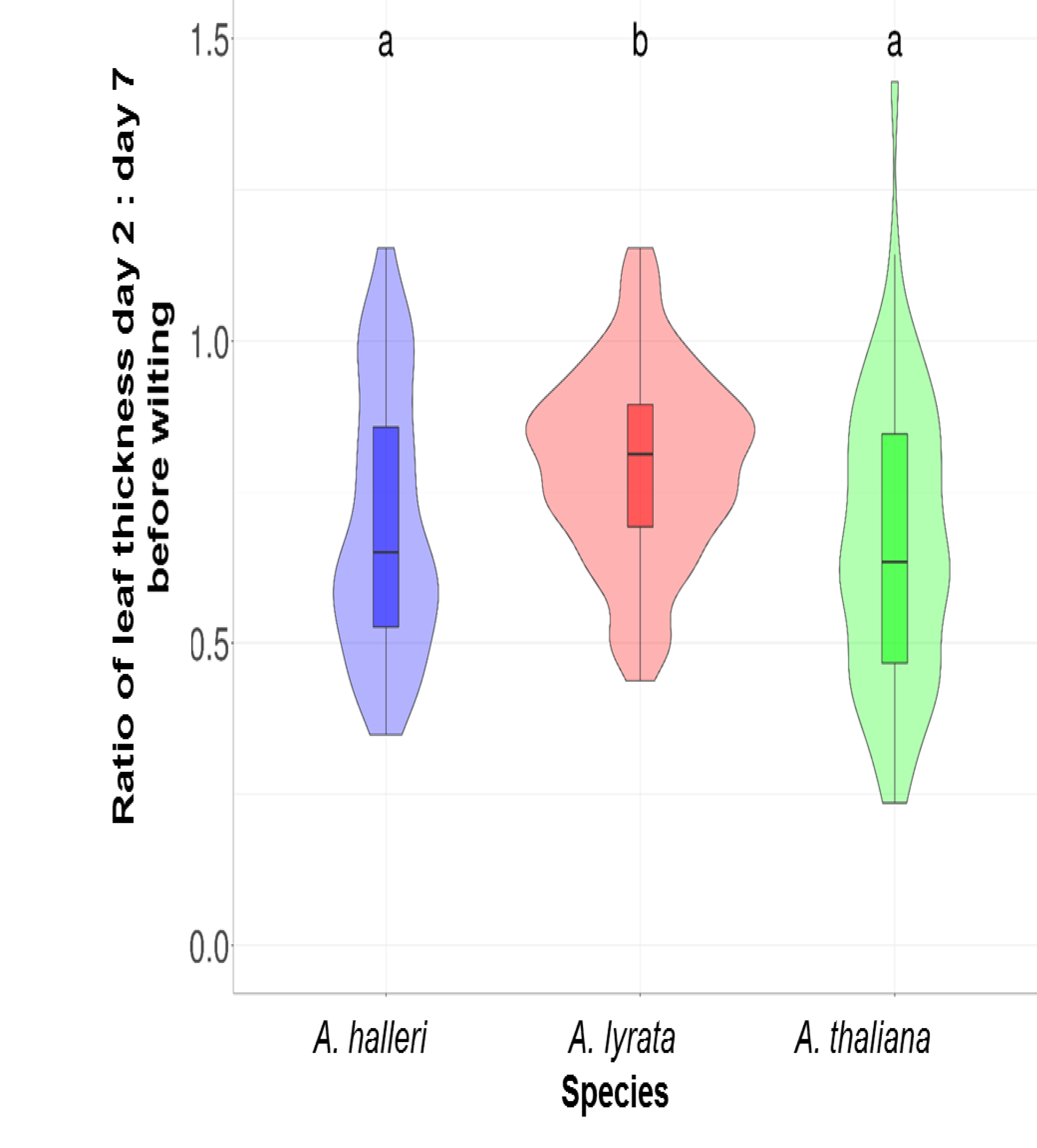
Relative leaf water loss seven days before wilting in *Arabidopsis halleri*, *A. lyrata*, and *A. thaliana*. This is equivalent to the ratio of leaf thickness at day two vs day seven before wilting. Boxplots with the same letter are not significantly different (*Tukey’s HSD, P value <0.05*). Results are shown for the second biological experiment.

### High photosynthesis efficiency in wilted A. halleri and A. lyrata plants

Photosynthesis efficiency was measured to evaluate the physiological status of plants at wilting. We used F_v_: F_m_ ratio, as indicator for the potential capacity of non-cyclic electron flow in the photosynthetic light reaction. Despite the collapsed or rolled leaves observed at wilting in *A. halleri* and *A. lyrata*, respectively, both still had a high photosynthetic capacity: on average 83 and 90%, respectively. By contrast, the photosynthetic capacity had significantly dropped in wilted *A. thaliana* rosettes (Supplementary Fig. S5, Supplementary Table S10).

### A. thaliana has the lowest survival rate

Individual plants were re-watered two days after observing symptoms of wilting. Two to three weeks after re-watering, we scored survival. The proportion of survivors was significantly lower in *A. thaliana* compared to *A. halleri* and *A. lyrata* (9% in *A. thaliana*, 85% in *A. halleri* and 84% in *A. lyrata*, Fig. 7, Supplementary Table S11). These differences were consistent across the two experiments (Supplementary Fig. S6).

**Figure 7:**
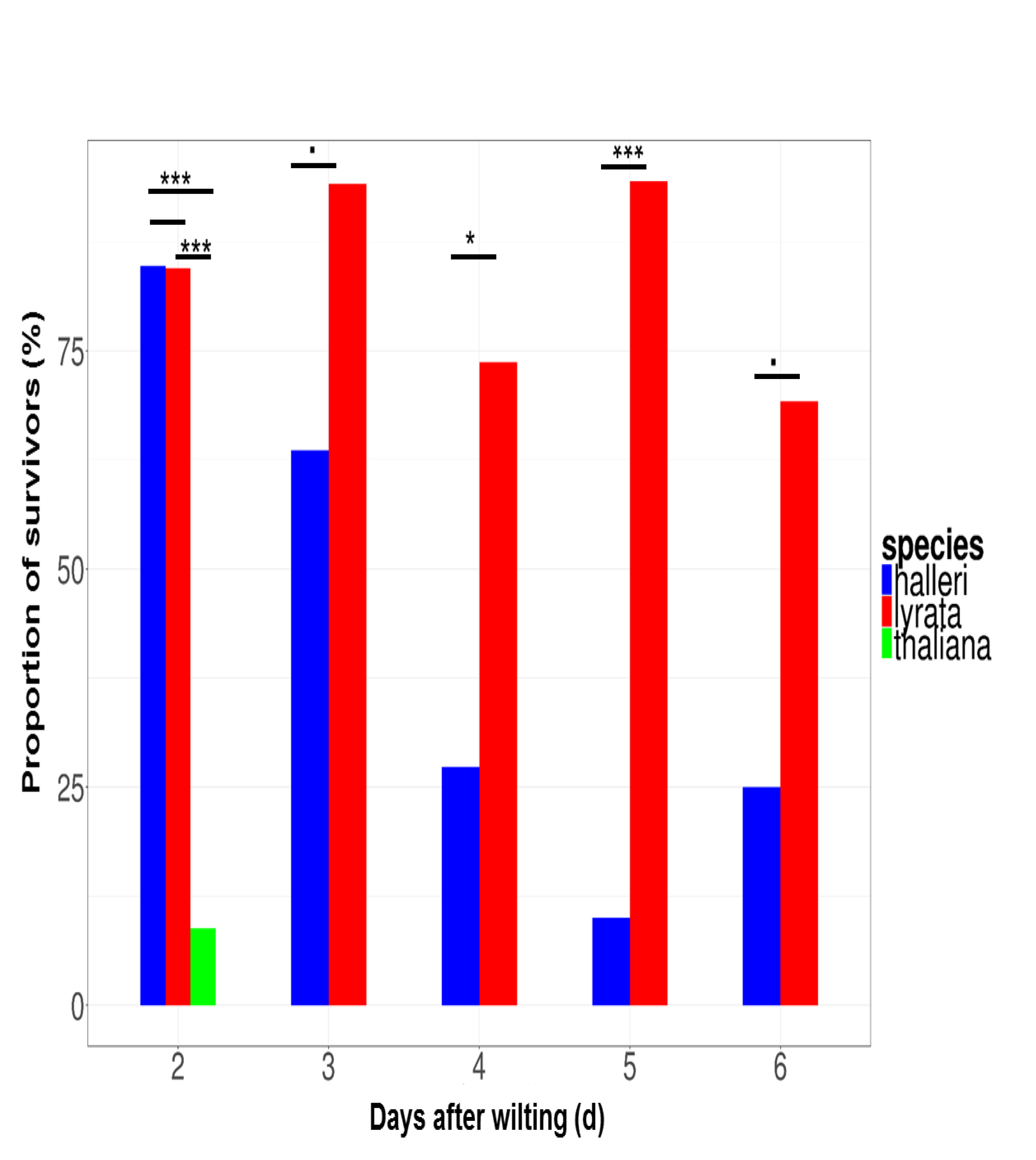
Average survival rate after re-watering following two to six days of wilting for *Arabidopsis halleri*, *A. lyrata*, and *A. thaliana*. Results are shown for the first biological replicate. Barplots with one star or more are significantly different (*Tukey’s HSD, Signif. codes:* ·, *P < 0.1;* *, *P < 0.05; **, P < 0.01; ***, P < 0.001; ns, not significant*).

To evaluate and compare the tolerance to wilting in *A. lyrata* and *A. halleri*, we ran an additional experiment examining whether extending the time from wilting to re-watering impacted survival. We detected a significant interaction effect of species and time to re-watering on survival (M4: Chi-Squared=234, DF= 1, DF residuals=252, P-value=1.615e^-04^). We observed that 70-85% of *A. lyrata* plants survived 3 to 6 day-long wilting periods (Fig. 7). In comparison, this percentage dropped to 10% for *A. halleri* plants after five days of wilting and this was significantly different between species (Fig. 7, M2: F_1, 26_= 20.681, P-value = 2.44e^-10^). These results indicate that *A. lyrata* is more tolerant to wilting than its sister species *A. halleri*.

### Efficient post-drought recovery in A. lyrata plants

We further, assessed the tolerance to wilting by comparing damage exhibited by plants that survived two days of wilting in *A. lyrata* and *A. halleri*. The interaction between species and the damage score was found to be significant (M4, F_3, 100_=2.96, P-value= 0.035). In *A. lyrata*, about 70% of plants showed a very low degree of damage in leaves, whereas in *A. halleri*, only 30% of plants had low damage levels (M4, Fig. 8, F_1, 25_= 24.063, P-value= 4.761e^-05^). We did not include *A. thaliana* in the statistical analysis because only 10 out of 60 plants survived wilting. These results confirmed that *A. lyrata* tolerates soil dehydration and wilting better than *A. halleri*.

**Figure 8:**
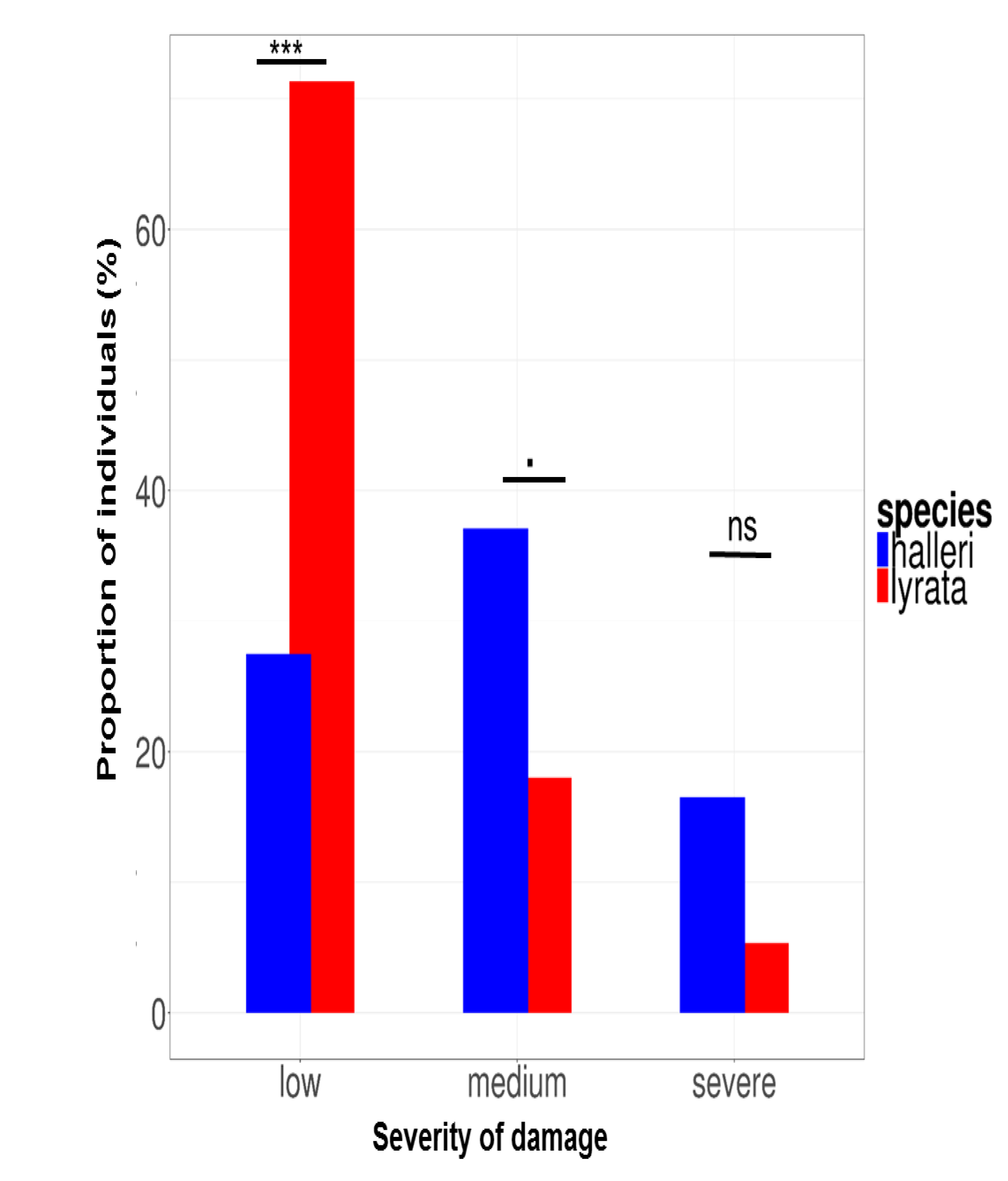
Damage scored on survivors to two days of wilting after resuming growth for *Arabidopsis halleri*, *A. lyrata*, and *A. thaliana*.. Results are shown for the second biological experiment. Barplots with one star or more are significantly different (*Tukey’s HSD, Signif. codes:* ·, *P < 0.1; *, P < 0.05; **, P < 0.01; ***, P < 0.001; ns, not significant*).

### Transcriptome analysis confirms that A. halleri is more sensitive to low SWC

*A. lyrata* and *A. halleri* both wilted at the same SWC but they differed in their survival following wilting. In order to gain insight into the molecular changes underpinning these differences, we performed a third dry-down experiment to collect leaf material in one representative accession of each of the sister species *A. halleri* and *A. lyrata* and examined the reaction to stress and recovery at the transcriptome level.

For each species, we compared transcript abundance at three time points during the dry-down experiment, i.e., at soil moisture 60%, soil moisture 20-25% and after recovery. The two species wilted at around 18% of soil moisture, as observed in the first two experiments, i.e., just below the soil moisture level at which leaf material was sampled. 107 and 976 genes changed expression level at 20-25 vs. 60% soil moisture in *A. lyrata* and *A. halleri*, respectively (FDR 0.1; fold-change >1.6). Only three genes were responsive in both species to the decrease in SWC and this was a random overlap (*hypergeometric test, P-value*=*0.99*3).

After recovery, 275 *A. lyrata* genes and 20 *A. halleri* genes had changed expression level compared to 60% SWC (Table 1). Since both species had similarly high survival rates upon two days of wilting and because new undamaged leaves were sampled, these differences are not due to survival differences. We conclude that *A. halleri* displayed a comparatively sharpened response to low SWC, whereas the transcriptome of *A. lyrata* was comparatively more altered after recovery.

**Table 1:** Number of significantly differentially expressed genes in *Arabidopsis halleri* and *A. lyrata* during the dry-down experiment at 20% of soil moisture or after recovery compared to expression before stress (60% of soil moisture).

|  |  | *A. halleri* | *A. lyrata* |
| --- | --- | --- | --- |
| **20% vs 60% of soil moisture** | Up | 253 | 36 |
|  | Down | 676 | 71 |
| **recovery vs 60% of soil moisture** | Up | 8 | 111 |
|  | Down | 12 | 156 |

In a previous study, 2975 and 5445 genes were shown to be responsive to two and 10 hours of dehydration in *A. thaliana* respectively (Matsui *et al.*, 2008). These drought-responsive genes were enriched in all sets of responsive genes identified in our study, either in *A. halleri* or in *A. lyrata*, at 20% soil moisture or after recovery (Table 2, *hypergeometric test*, maximum p ≤ 8.77E-19). This confirmed that our protocol succeeded in activating dehydration responsive genes. The list of significantly differentially expressed genes (including only AGI codes) is provided in Supplementary Table S12.

**Table 2:**
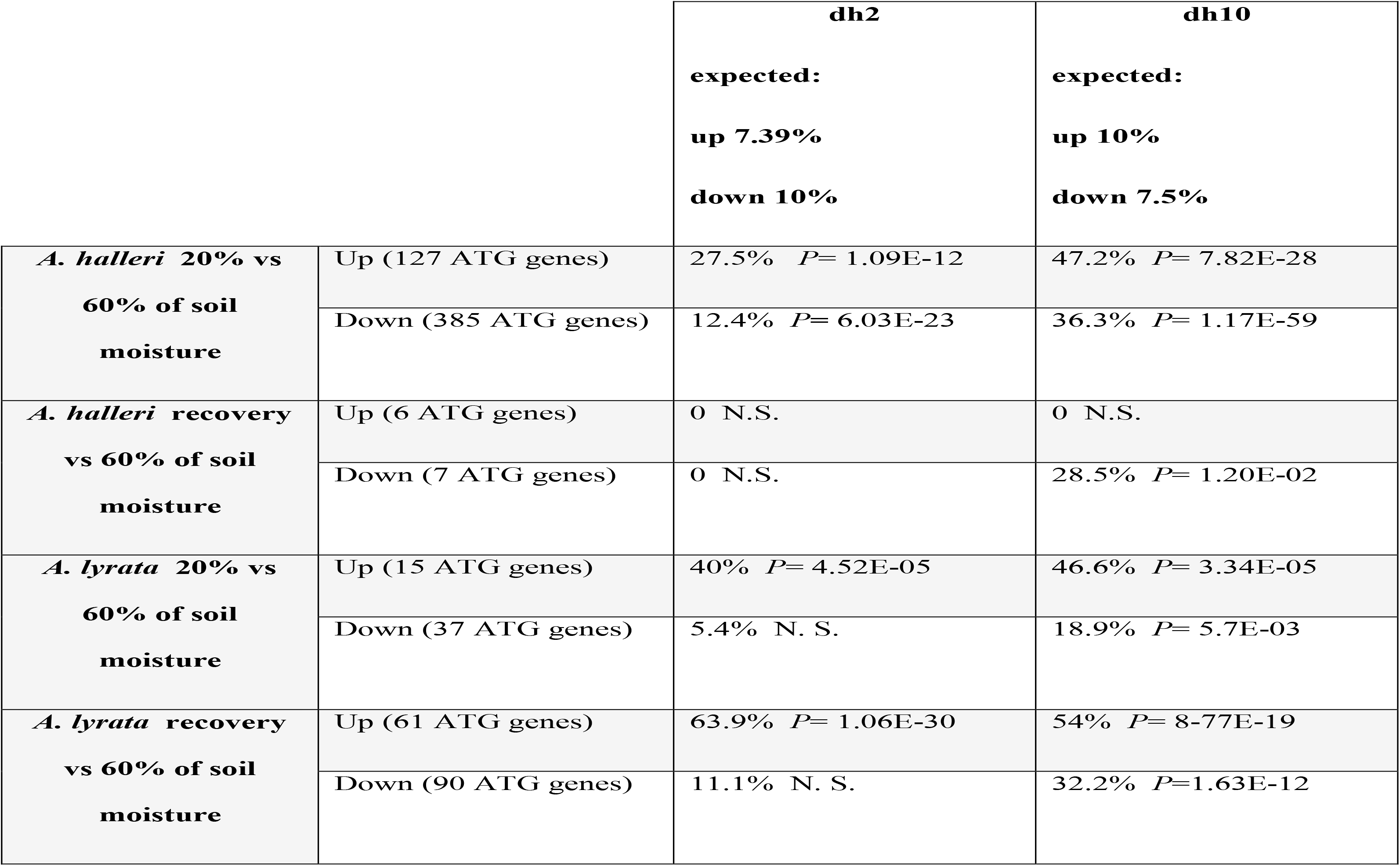
Percentage of differentially expressed genes that overlap with differentially expressed genes reported in Matsui *et al.*, (2008)after 2 h (dh2) and 10 h (dh10) of dehydration stress (N.S.: not significant). The random expectation of overlap % is indicated in bold on the top row.

|  |  | dh2 | dh10 |
| --- | --- | --- | --- |
|  |  | expected:<br>up 7.39%<br>down 10% | expected:<br>up 10%<br>down 7.5% |
| ***A. halleri* 20% vs**<br>**60% of soil**<br>**moisture** | Up (127 ATG genes) | 27.5% $P= 1.09\text{E-}12$ | 47.2% $P= 7.82\text{E-}28$ |
| | Down (385 ATG genes) | 12.4% $P= 6.03\text{E-}23$ | 36.3% $P= 1.17\text{E-}59$ |
| ***A. halleri* recovery**<br>**vs 60% of soil**<br>**moisture** | Up (6 ATG genes) | 0 N.S. | 0 N.S. |
| | Down (7 ATG genes) | 0 N.S. | 28.5% $P= 1.20\text{E-}02$ |
| ***A. lyrata* 20% vs**<br>**60% of soil**<br>**moisture** | Up (15 ATG genes) | 40% $P= 4.52\text{E-}05$ | 46.6% $P= 3.34\text{E-}05$ |
| | Down (37 ATG genes) | 5.4% N. S. | 18.9% $P= 5.7\text{E-}03$ |
| ***A. lyrata* recovery**<br>**vs 60% of soil**<br>**moisture** | Up (61 ATG genes) | 63.9% $P= 1.06\text{E-}30$ | 54% $P= 8.77\text{E-}19$ |
| | Down (90 ATG genes) | 11.1% N. S. | 32.2% $P= 1.63\text{E-}12$ |

### Different GO categories are regulated in the two species

Analysis of enrichment in Gene Ontology (GO) categories confirmed that different sets of genes were activated in the two species at each sampling stage. In *A. halleri* many genes involved in growth and development were down regulated when SWC decreased to 20-25%, (Table 3). These functions were not enriched in *A. lyrata* samples collected at the same time, instead genes involved in response to water deprivation and in ethylene and ABA signaling pathways were up regulated in *A. lyrata* after recovery (Table 3). Several GO terms appeared enriched, including isopentenyl diphosphate metabolic process, response to water deprivation, hyperosmotic salinity response, photosynthesis light reaction, response to chitin, photosystem II assembly, and maltose metabolic process (Table 3). They were also enriched among genes responding to mild drought stress in *A. thaliana*, although the direction of the gene expression change was not the same (Des Marais *et al.*, 2012). We further observed that genes with altered expression in *A. halleri* were enriched for genes functioning in plastid organization, pentose-phosphate shunt and photosystem II assembly. These three GO categories harbor an excess of *cis*-acting changes in the *A. halleri* lineage in response to dehydration stress (He *et al.*, 2016).

**Table 3:** GO Categories Showing a Significant Enrichment (P < 0.01) among differentially expressed genes between 20% and 60% of soil moisture and between recovery and 60% of soil moisture for *Arabidopsis halleri* and *A. lyrata*.

|  | GO.ID | Term | pvalue | Gene regulation |
| --- | --- | --- | --- | --- |
| *A. halleri*<br>20% vs<br>60% of<br>soil<br>moisture | GO:0015979 | photosynthesis | 0.0011 | down |
|  | GO:1901576 | organic substance biosynthetic process | 0.0013 | down |
|  | GO:0044711 | single-organism biosynthetic process | 0.0014 | down |
|  | GO:0051188 | cofactor biosynthetic process | 0.0023 | down |
|  | GO:0008283 | cell proliferation | 0.0035 | down |
|  | GO:0006098 | pentose-phosphate shunt | 0.0041 | down |
|  | GO:0009965 | leaf morphogenesis | 0.0048 | down |
|  | GO:0009657 | plastid organization | 0.0059 | down |
|  | GO:0042254 | ribosome biogenesis | 0.0059 | down |
|  | GO:0006084 | acetyl-CoA metabolic process | 0.0064 | down |
| *A. lyrata*<br>recovery<br>vs 60% of<br>soil<br>moisture | GO:0006098 | pentose-phosphate shunt | 0.000043 | down |
|  | GO:0010200 | response to chitin | 0.000051 | up |
|  | GO:0010207 | photosystem II assembly | 0.00007 | down |
|  | GO:0000023 | maltose metabolic process | 0.00017 | down |
|  | <u>GO:0009873</u> | ethylene-activated signaling pathway | 0.0002 | up |
|  | GO:0019252 | starch biosynthetic process | 0.00039 | down |
|  | GO:0009612 | response to mechanical stimulus | 0.0015 | up |
|  | GO:0009414 | response to water deprivation | 0.0029 | up |
|  | GO:0042538 | hyperosmotic salinity response | 0.0043 | up |
|  | GO:0051707 | response to other organism | 0.005 | up |
|  | GO:0009657 | plastid organization | 0.00571 | down |
|  | GO:0050790 | regulation of catalytic activity | 0.00763 | down |
|  | GO:0042742 | defense response to bacterium | 0.00784 | down |
|  | GO:0009738 | abscisic acid-activated signaling pathway | 0.0086 | up |

## DISCUSSION

In our experimental design, we have used several accessions per species as we were interested in comparing the drought stress response of the three related species, while accounting for variation within species. To exclude the possibility that our results are influenced by a previous history of stress, we discarded sick or slow growing plants and studied the drought response of vigorously growing individuals. Our results showed genotypic differences in initial leaf thickness, initial stomatal density or initial rosette area, but the response to depletion in SWC did not reveal significant differences between accessions. Differences in the response to water depletion therefore revealed fixed interspecific differences in avoidance and tolerance strategies to drought stress.

### Critical SWC does not reflect ecological differences between A. halleri and A. lyrata

The sister species *A. lyrata* and *A. halleri* have separated recently and gene flow between the clades is still detectable (Novikova *et al.*, 2016). Yet, the two species display marked differences in ecological preference (Clauss & Koch, 2006). Ellenberg indices, which are reliable estimates of ecological preferences in Central Europe, show that *A. lyrata* is found in very dry areas with a soil humidity index (F) of 3, while *A. halleri* occurs in habitats where water is less limiting (F= 6) (Ellenberg & Leuschner, 2010). We were therefore surprised to observe that *A. halleri* and *A. lyrata* individuals wilted at identical SWC. In addition, contrary to our expectations, the ruderal species *A. thaliana* tolerated markedly lower SWC than its perennial relatives. Altogether, these observations show that the ecological preferences of *A. lyrata*, *A. halleri* and *A. thaliana* are not explained by the SWC threshold at which wilting symptoms appear.

### *A. halleri* is directly exposed to stress caused by low SWC

We observed that *A. halleri* was the fastest to consume the water contained in the soil. In pots where *A. halleri* individuals grew, SWC decreased significantly faster (Supplementary Fig. S3). *A. halleri* also displayed the strongest correlation between plant size and the rate of water consumption and an accelerated decrease in leaf thickness preceding the onset of wilting (Fig. 4-6). At 25% soil water content, i.e. shortly before the appearance of the first wilting symptoms, the rate of decrease in leaf thickness accelerated in *A. halleri* compared to *A. lyrata*. This turning point coincided with a change in the expression levels of a larger number of genes belonging to stress-repressed GO categories such as leaf morphogenesis, cell proliferation, or photosynthesis. The down-regulation of growth-related genes we observed, even before wilting symptoms appear, indicates that the plant experiences direct stress at the cellular level as SWC approaches the limiting threshold. In agreement with the high levels of stress it experienced, *A. halleri* also showed a comparatively higher damage when survivors resumed growth after stress.

Although less tolerant to wilting than *A. lyrata*, *A. halleri* did display some level of tolerance, because it was comparatively more tolerant than *A. thaliana* as it did survive two days of wilting. Yet, of the three species, *A. halleri* clearly displayed the weakest levels of drought avoidance, being almost directly exposed to stress caused by decreasing SWC. *A. halleri* thrives in more competitive habitats than its relatives (Clauss & Koch, 2016; Stein *et al.*, 2017), and competitive ability generally evolves in a trade-off with stress tolerance in plant species (Grime *et al.*, 1977; Sreenivasulu *et al.*, 2012). It is therefore possible that improved competitive ability was selected in this lineage at the expense of tolerance and avoidance mechanisms. Such evolutionary scenarios have been documented in several grass species (Fernández & Reynolds, 2000; Liancourt *et al.*, 2005; Sugiyama, 2006). Interestingly, we have previously observed that an excess of *cis*-acting changes up-regulating gene expression after one hour of dehydration had accumulated in the *A. halleri* lineage in several functions that the more tolerant species *A. lyrata* down-regulates during recovery (He *et al.*, 2016). It is therefore possible that the decrease in tolerance and avoidance of drought stress was advantageous in the context of selection for increased competitive ability.

### A. lyrata displays avoidance and tolerance responses to soil dehydration

By comparison with *A. halleri*, *A. lyrata* displayed a more parsimonious use of water. *A. lyrata* plants displayed both a lower rate of water consumption and markedly lower damage levels after resuming growth. In addition, we observed that *A. lyrata* plants had the ability to survive longer durations of wilting than both *A. halleri* and *A. thaliana* (Fig. 7). It is also the only species that showed adaxial leaf rolling, a phenotype favoring drought avoidance in plants (Oppenheimer, 1960; O’Toole & Moya, 1978; Jones, 1979; Clarke, 1986). Leaf rolling indeed reduces transpiration rate by reducing the effective leaf area. Altogether, this indicates that exposure to limiting SWC is comparably less damaging in *A. lyrata*.

The transcriptome response to decreasing SWC corroborated this observation, by documenting lower levels of cellular stress in *A. lyrata* immediately before wilting, compared to *A. halleri*. Only a few genes changed expression before wilting in *A. lyrata*. We further observed that among genes down-regulated after recovery, the most enriched GO category is ‘pentose-phosphate shunt’ (p<5.10^-5^), a metabolic pathway involved in the scavenging of reactive oxygen intermediates that is strongly activated by abiotic stress (Mittler, 2002; Kruger & von Schaewen, 2003). Several additional GO functions associated with drought stress, such as ‘hyperosmotic salinity response’, ‘response to water deprivation’, ‘abscisic acid-activated signaling pathway’, ‘ethylene-activated signaling pathway’, and ‘response to chitin’ were up-regulated in *A. lyrata* during recovery. The latter functions seem to have a dynamic role in drought stress. In *A. thaliana*, they were up-regulated by severe fast wilting (Matsui et al. 2008) but down-regulated by mild stress (Des Marais *et al.*, 2012). Their up-regulation after recovery in *A. lyrata*, in the absence of obvious stress, shows that the reaction of this species to lowering SWC contrasts not only with that displayed by *A. halleri* but also with that known for *A. thaliana*. The absence of a strong modification of the expression of drought-stress responsive genes at SWC approaching critical levels in *A. lyrata*, combined with a high survival rate, further indicates that this species has the ability to i) minimize its exposure to the stressful consequences of low soil water content and ii) maximize its ability to survive severe dehydration. It thus deploys both avoidance and tolerance strategies.

Whether the lower stomata density observed in *A. lyrata* (Fig. 1a) contributes to its improved ability to cope with limiting water availability is difficult to evaluate with our data. Indeed, increased stomata density has been associated with decreased WUE both within and between species (Carlson, Adams, & Holsinger, 2016; Muchow & Sinclair, 1989; Reich, 1984; Anderson & Briske, 1990; Pearce, Millard, Bray, & Rood, 2006; Doheny-Adams *et al.*, 2012; Liu *et al.*, 2012). Yet, in monkey flowers and in *Arabidopsis thaliana*, lower stomatal density was associated with higher WUE (Wu *et al.*, 2010,Dittberner et al. 2018). The consequences of modification in stomata density and size on the plant’s ability to cope with limiting water supply are, in fact, not easily predictable. First, water use efficiency can increase as a result of either increased stomata density or increased stomata size because larger stomata close more slowly (Raven, 2014). Second, the two traits generally correlate negatively (Dittberner et al. 2018, Hetherington and Woodward, 2003). Third, parameters independent of stomata patterning such as photosynthetic ability can also contribute to variation in WUE, as reported recently in *A. thaliana* (Dittberner et al. 2018,Farquhar et al. 1989). Fourth, stomata patterning changes in *A. lyrata* plants when exposed to limiting water supply (Paccard et al. 2014). Our data reveals that in well-watered greenhouse conditions *A. lyrata* did not show a globally higher WUE than *A. halleri* (Fig. 1b), despite significant differences in stomata density and size. Future work will have to investigate the impact of modifications in stomata patterning on interspecific differences in tolerance and avoidance in the face of limiting SWC.

### High levels of stress avoidance associate with low tolerance to drought in A. thaliana

In annual species, seasonal drought can be a potent source of selection for accelerated flowering and faster cycling (Franks *et al.*, 2007; Fitter & Fitter, 2002). *A. thaliana* was also expected to maximize its resource investment into growth and reproduction and show a lower level of stress tolerance compared to its perennial relatives. Here, we focused on late flowering *A. thaliana* accessions that in the conditions we imposed could not accelerate their development to escape drought. Thus, we cannot conclude on the relative investment of Arabidopsis species in escape strategies, but our experimental set up allowed an interspecific assessment of avoidance and tolerance to wilting. Contrary to expectations, we observed that our sample of accessions could persist at lower SWC than both of their perennial relatives, *A. lyrata* and *A. halleri* (Fig. 3A). In addition, the delayed decrease in leaf thickness observed in *A. thaliana* shows that, compared to the other two species, it is able to maintain its leaf water content at lower SWC (Fig. 5). This therefore suggests that the annual species *A. thaliana* also employs stress avoidance mechanisms. The ability of this annual species to escape stress by accelerating development has therefore not led to the loss of mechanisms favoring the maintenance of internal water potentials. Indeed, the production of proline, which is both an osmoprotectant and an anti-oxidant, δ^13^C, a proxy measuring WUE, as well as the maintenance of photosynthesis during terminal drought have been documented to play a role in local adaptation in this species (Verslues & Juenger, 2011; Kesari *et al.*, 2012; Exposito-Alonso *et al.*, 2017; Dittberner *et al.*, 2018).

*A. thaliana*, however, was not able to tolerate wilting. We observed a marked decrease in the photosynthetic capacity at wilting in this species, as previously reported in several species such as *Hordeum vulgare*, *Hibiscus rosa-sinensis*, and *Andropogon gerardii* (Golding & Johnson, 2003; Muñoz & Quiles, 2013; Maricle *et al.*, 2017). In addition, *A. thaliana* did not survive after two days of wilting, although its perennial relatives displayed markedly higher survival rates. The annual species therefore appears to have evolved lower levels of tolerance to wilting.

We detected no significant variation for the response to decreasing SWC between the *A. thaliana* accessions included in this study, however, we cannot conclude that the avoidance capacity and the low tolerance to wilting we observed is fixed in the species. The *A. thaliana* population we used consisted of a set of late-flowering accessions from Spain that could not accelerate flowering fast enough to escape stress. This set of accessions is not necessarily representative of the whole species. *A. thaliana* is broadly distributed and its accessions can form ecotypes with markedly different levels of stress resistance (May *et al.*, 2017). Furthermore, two recent studies indicate that Swedish accessions have a comparatively greater capacity to face dry conditions, probably because the short favorable season of Scandinavia constrains them to face limiting water availability when it strikes (Exposito-Alonso *et al.*, 2017,Dittberner *et al.*, 2018).

This study documents the contrasting reactions deployed by *Arabidopsis* species in response to lowering SWC. In the face of their respective ecologies, these diversified reactions likely reflect the priority shifts imposed by divergent ecologies and life cycles. Future studies aiming at dissecting the genetic and molecular underpinning of these differences promise to teach us much about the processes accompanying ecological diversification in plant species.

## ACKNOWLEDGEMENTS

We thank Agustín Arce, Kim Steige, Ulrike Goebel, Hannes Dittberner, Veronica Preite, Julia Plewka, Nina Grisard, Martin Rippin, and Kirsten Bell for their technical support. Ute Krämer for providing *A. halleri* accessions, commenting on experimental set up, and commenting on the manuscript draft. The Cologne Center for Genomics performed HiSeq4000 RNA sequencing. The phenotypic data is available as supplementary information (Suppl. Table 13).

## Supporting Information

**Figure S1:** Summary of short read mapping to the *A. lyrata* reference genome V1.

**Figure S2:** Wilting day and soil moisture at wilting for the two first biological experiments of the drying-down experiments.

**Figure S3:** Soil water content during the first 7 days after water withdrawal.

**Figure S4:** Initial rosette area and leaf thickness of the plants used in the second biological experiments of the drying-down experiment.

**Figure S5:** Photosynthesis efficiency at wilting.

**Figure S6:** Proportion of surviving *A. halleri*, *A lyrata*, and *A. thaliana* plants 2 days after re-watering for the two first biological experiments.

**Table S1**: List of accessions used for the dry-down experiments.

**Table S2:** Phenotypes measured in the three drying-down experiments.

**Table S3:** Number of accessions used in the three drying-down experiments.

**Table S4**: Summary statistics of the multiple comparison of the wilting day between species.

**Table S5**: Summary statistics of the multiple comparison of the soil moisture at wilting between species.

**Table S6**: Summary statistics of the multiple comparison of the initial rosette area between species.

**Table S7**: Summary statistics of the multiple comparison of the initial leaf thickness between species.

**Table S8**: Summary statistics of the multiple comparison of the relative leaf water loss 7 days before wilting between species.

**Table S9**: Summary statistics of glm testing the effect of interaction between species and desiccation rate on the relative loss of leaf water content before wilting.

**Table S10**: Summary statistics of the multiple comparison of the photosynthetic efficiency at wilting between species.

**Table S11**: Summary statistics of the multiple comparison of the survival rate 2 days after re-watering between species.

**Table S12:** Differentially expressed genes identified for each of *Arabidopsis halleri* and *A. lyrata* between 20 and 60% of soil moisture and bewteen recovery and 60% of soil moisture.

**Table S13:** Phenotypic data collected in this study. See methods for details on the measurements and experimental procedures.

